# Mesenchymal stromal cells affect CD8 naïve to memory subsets polarization by down-modulating IL12Rβ1 and IL2Rα signaling pathways

**DOI:** 10.1101/2023.02.28.530378

**Authors:** Andrea Papait, Elsa Vertua, Patrizia Bonassi Signoroni, Anna Cargnoni, Marta Magatti, Francesca Romana Stefani, Jacopo Romoli, Antonietta Rosa Silini, Ornella Parolini

## Abstract

Adaptive immunity is typified by specificity and memory. Immune memory protects from subsequent infection and relies on functional, fully activated, T lymphocytes. Mesenchymal stromal cells (MSC) have been investigated for their potential therapeutic applications in a variety of diseases in which a dysregulated immune response plays a central role in pathogenesis or progression. Notably, for adaptive immunity, MSC can reduce the activation and cytotoxic activity of CD8^+^ T lymphocytes as well as the polarization of CD4^+^ T lymphocytes toward inflammatory subsets while favoring polarization toward the T regulatory subset. Although MSC have been widely reported to impact CD4 T cell proliferation and polarization, how MSC affect T-cell commitment toward memory subsets is still not known. Here, we report for the first time that MSC isolated from the amniotic membrane of human term placenta (hAMSC) determine T cell fate. We show that hAMSC influence naïve CD8^+^ T cell activation and differentiation by downregulating mTOR pathway activation and modulating the expression of Tbet and Eomes, master regulators of the commitment of naïve CD8^+^ T cells toward memory precursor effector cells (MPECs). This effect can be partly attribute to the ability of hAMSC to reduce the phosphorylation of STAT4 and STAT5, two transcriptional factors downstream IL-12Rβ1 and IL-2Rα receptors. Our results unravel a novel feature of MSC, offering new mechanistic insights into the effects of MSC in the treatment of diseases characterized by an altered activation of memory subsets, such as autoimmune diseases and graft versus host disease.

## Introduction

The formation of an adaptive T cell memory repertoire is a key feature of the primary adaptive immune response.

Shortly after activation, CD8 effector cells are differentiated into short-lived effector cells (SLECs), characterized by low survival and high proliferation potential, and long-lived memory precursor effector cells (MPECs), characterized by developing into memory cells (*1, 2*).

Antigen-specific memory CD8 T cells constitute the long-lived arm of immunity that rapidly protects the organism from secondary infections or tumors (*3*), and the inability to establish proper CD8-driven immunological memory underlies numerous pathologies such as autoimmune diseases and allograft rejection (*4-6*).

Mesenchymal stromal cells (MSCs) are versatile stromal components whose known role in modulating immunity and inflammation has been exploited in numerous clinical trials ranging from their use in the treatment or prevention of graft versus host disease (GvHD) to the treatment of autoimmune diseases such as Crohn’s disease and neurodegenerative disorders (*7, 8*). MSCs isolated from the amniotic membrane of the human placenta (hAMSCs) have been shown to influence several components of adaptive immunity, including T cells. Their ability to perform immune-related functions relies on several mechanisms such as suppression of T lymphocyte activation (*9, 10*), blocking polarization toward inflammatory Th subsets (*10-12*), and induction of Treg regulatory cells (*11, 12*). In addition, they have been shown to block the maturation and differentiation of monocytes into antigen-presenting cells (APCs) while promoting polarization and the acquisition of anti-inflammatory properties typical of M2 macrophages (*13, 14*). Preclinical studies have shown that hAMSCs can exert therapeutic effects in animal models of acute and chronic inflammation-related diseases (*15-19*). Furthermore, it is now known that many of the immunoregulatory actions of hAMSCs occur via paracrine mechanisms, underscoring the clinical relevance of the factors contained and mediated in their secretome (20-22).

Here, we investigated a topic not yet explored, namely the potential impact of hAMSC and their secreted factors on the maturation and formation of the memory CD8 lymphocyte repertoire.

Our study focused on the early post-activation stages of naïve CD8 lymphocytes, in which the expression of Tbet and Eomesodermin (Eomes), master regulators of T cell function, drive the differentiation of naïve CD8 T cells into SLECs and MPECs, respectively. We discovered a novel immunological property of hAMSC targeting the formation of memory CD8 T lymphocytes based on the control of naïve CD8 cell to MPEC polarization via Tbet and Eomes and the ability of hAMSC to downregulate the phosphorylation of AKT and mTOR. Importantly, the immunomodulatory capacity of hAMSC may be correlated with its ability to modulate the expression and activation of two receptors, IL -2 receptor alpha (IL -2Rα) and IL -12 receptor beta 1 (IL -12Rβ1), which is reflected in the downstream downregulation of STAT5 and STAT4, two important transcription factors involved in the differentiation of naïve CD8 lymphocytes.

## Results

### hAMSC harbor the potential to determine T cell fate

To investigate the impact of hAMSC on immunological CD8 T cell memory commitment, we first analyzed the effect of hAMSC on the activation and engagement of CD8 T lymphocytes within the PBMC stimulated with anti CD3 mAbs over time.

Co-culture of PBMCs with hAMSC reduced proliferation of both naïve and differentiated CD8 T cell subsets (central memory (CM), effector memory (EM), and effector memory RA (TEMRA)) at day 3 (Fig. 1A). This effect was no longer detectable at day 7, at which time CD8 proliferation was drastically reduced and the effect of hAMSC on CD8 proliferation was difficult to assess (Fig. 1D). To determine whether the effect on T-cell proliferation reflected changes in CD8 T-cell memory commitment, we analyzed the proportion of different memory subsets after treatment with hAMSC. At day 3, differentiation of naïve cells in CM, EM, and TEMRA was not affected by hAMSC conditioning (Fig. 1B). Conversely, at day 7, the EM pool was reduced and its reduction was counterbalanced by maintaining a pool of naïve and CM cells (Fig. 1E). Accordingly, flow cytometry analysis of Tbet and Eomes, two transcription factors involved in the early stages of CD8 lymphocyte commitment to memory subsets, showed a drastic reduction in the expression of these two transcription factors after co-culture with hAMSC. This effect was observedat day 3 (Fig. 1C) and became even more prominent at day 7 (Fig. 1F).

**Fig. 1.**
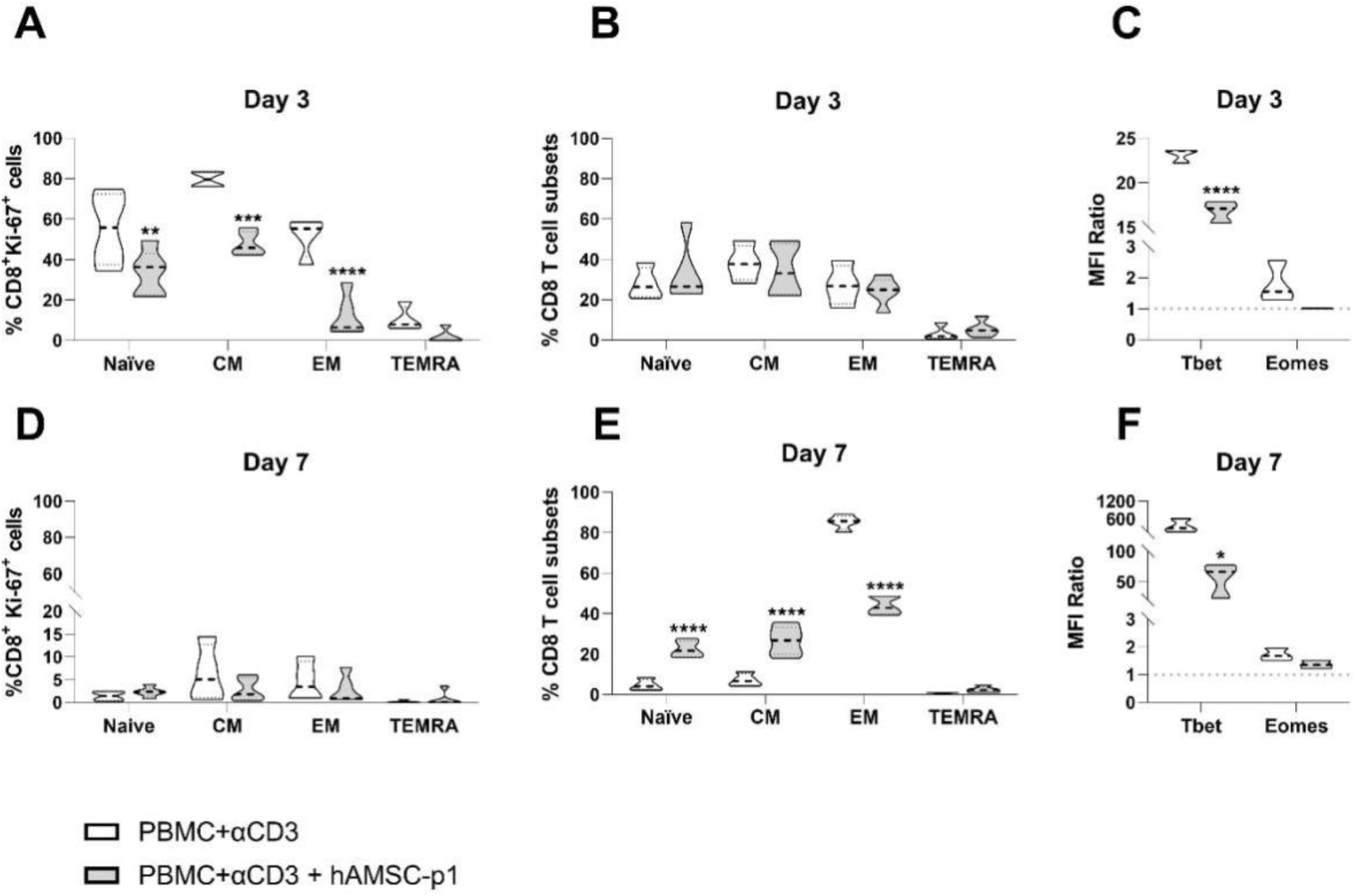
hAMSC regulate CD8 T cell fate within the PBMC pool. PBMC stimulated with antiCD3 antibody were co-cultured with hAMSC p1 (gray) for 3 (upper panels) or 7 (lower panels) days. Cells were then analyzed by flow cytometry to discriminate the CD8 naïve T cell pool from the memory subsets as follows: naïve T lymphocytes (CCR7+CD45RO-), central memory (CM) (CCR7+CD45RO+), effector memory (EM) (CCR7−CD45RO-) and effector memory RA (TEMRA) (CCR7−CD45RO-). Cell proliferation was evaluated as percentage of Ki67+ cells for the different CD8+ subsets investigated at day 3 (A) and 7 (D). CD8 T cell differentiation is represented as the percentage of the different subsets identified at day 3 (B) and 7 (E). Expression of the transcription factors Tbet and Eomes is represented as median fluorescence intensity (MFI) ratio evaluated at day 3 (C) and 7 (F). Results are displayed as violin plots showing median (dashed line), 25th and 75th quartiles (*p < 0.05, **p < 0.01, ****p < 0.001); stimulated PBMC alone represent the control group (white); N ≥ 3 independent experiments.

### hAMSC directly orchestrate T lymphocyte commitment

To define the direct role of hAMSC in CD8 T cell commitment, experiments were also performed with purified CD8 naïve T lymphocytes stimulated with antiCD3 and antiCD28 mAbs in the presence of IL -12 and IL -2 (*23, 24*).

Activated naïve CD8 T cells proliferated and progressively committed towards the subtypes CM and EM. This process was reversed by co-culture with hAMSC. Indeed, cell proliferation began to decline at day 3, became increasingly evident at day 7, and persisted until day 10. Consequently, cell commitment was impaired, with retention of the naïve fraction being very evident at day 7 and reflected in a parallel decrease in polarization toward the CM and EM subpopulations (Figs. 2A-B). To visualize the differentiation process over time and to understand whether hAMSC could induce the acquisition of distinctive features in CD8 T cells and/or differentiation into a new cellular subset, we performed high-dimensional flow cytometry analysis. We performed unsupervised analyzes on naïve CD8 lymphocytes to investigate cell fate development at different time points (day 3, 7, and 10). We employed several methods for multidimensional analysis, including FlowSOM meta-clustering in conjunction with a dimensionality reduction method such as Uniform Manifold Approximation and Projection (UMAP) (Fig. 2C). Finally, we identified specific subclusters using the Cluster Explorer plugin. In these 3D analyzes, we specifically examined the effects on CD183, a marker that is rapidly induced on naïve T lymphocytes after activation (*25*), along with the expression of the proliferation marker Ki67 and identified 8 clusters. We then reduced data complexity by pooling the closely spaced clusters and dividing the cells into 6 subgroups identified as follows: Naïve Ki67-, Naïve Ki67+, CM Ki67-, CM Ki67+, EM Ki67-, and EM Ki67+ (Fig. 2C).

**Fig. 2.**
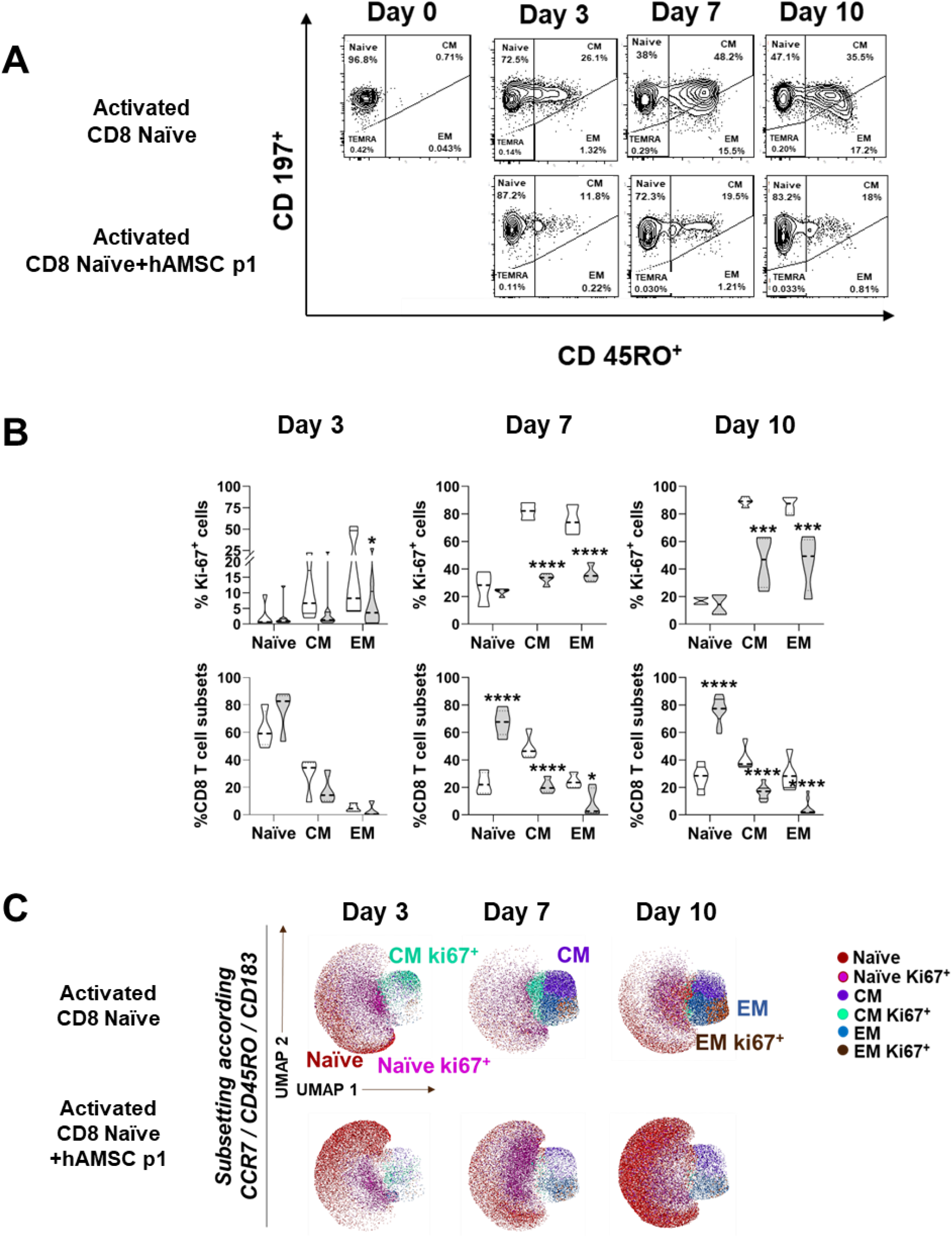
hAMSC directly orchestrate T lymphocyte commitment. Purified CD8 naïve T lymphocytes were stimulated with antiCD3, antiCD28 and the exogeneous administration of IL12 and IL2, and cultured in the presence of hAMSC (gray). (A-B) CD8 T cells were allowed to differentiate for 10 days and the degree of proliferation (as percentage of Ki67+ cells) and commitment (as percentage of the different T cell subsets) evaluated at day 3, 7 and 10 by flow cytometry. CD8 T cells were distinguished based on the differential expression of CD197 and CD45RO in naïve (CD45RO-CD197+), central memory (CM) (CD197+CD45RO+) and effector memory (EM) (CD45RO+CD197-). (C) Uniform Manifold Approximation and Projection (UMAP) representation of the CD8+ T cell landscape obtained by Clusterexplorer plugin. Cells were stratified for CCR7, CD45RO and CD183 and subsequently clustered. Results in (A) and (B) are displayed as violin plots showing median (dashed line), 25th and 75th quartiles (*p < 0.05, **p < 0.01, ****p < 0.001); stimulated CD8 T cells alone represent the control group (white); N ≥ 3 independent experiments.

### hAMSCs influence early phases of CD8 naïve T cell commitment

To further evaluate hAMSC as determinants of T cell fate, we analyzed their ability to drive the differentiation of naïve CD8 T cells toward the SLEC and MPEC subtypes. These two subtypes can be distinguished by the differential expression of two surface markers, CD183 and KLRG1. SLECs are defined as CD183-KLRG1+, whereas MPECs are characterized by expression of CD183 and loss of positivity for KLRG1. We also examined the frequency of early effector cells (EECs), which function as master precursor effector T cells and are characterized by co-expression of the two markers mentioned above (CD183+KLRG1+) (*26*), and the frequency of double-negative effector cells (DNECs), whose role in CD8 lymphocyte commitment remains unclear (*26*).

Having confirmed the ability of activated naïve CD8 T lymphocytes to differentiate into MPECs in the presence of exogenous IL -12 and IL-2 (Fig. 3A) (43), we next examined the effects of hAMSC on this early commitment phase. As shown in Fig. 3A, treatment with hAMSC resulted in a reduction in the MPEC compartment in terms of both relative frequency and absolute number. In parallel, the percentage of DNEC fraction increased, whereas no difference was observed in the total amount of CD8 T cells. The transcription factors Tbet and Eomes promote the differentiation of naïve CD8 lymphocytes (*27*). Here, we observed a substantial reduction in the MFI ratio of Tbet and Eomes at day 3 from challenge with hAMSC, a reduction that persisted until day 10, although not significantly (Fig. 3B).

**Fig. 3.**
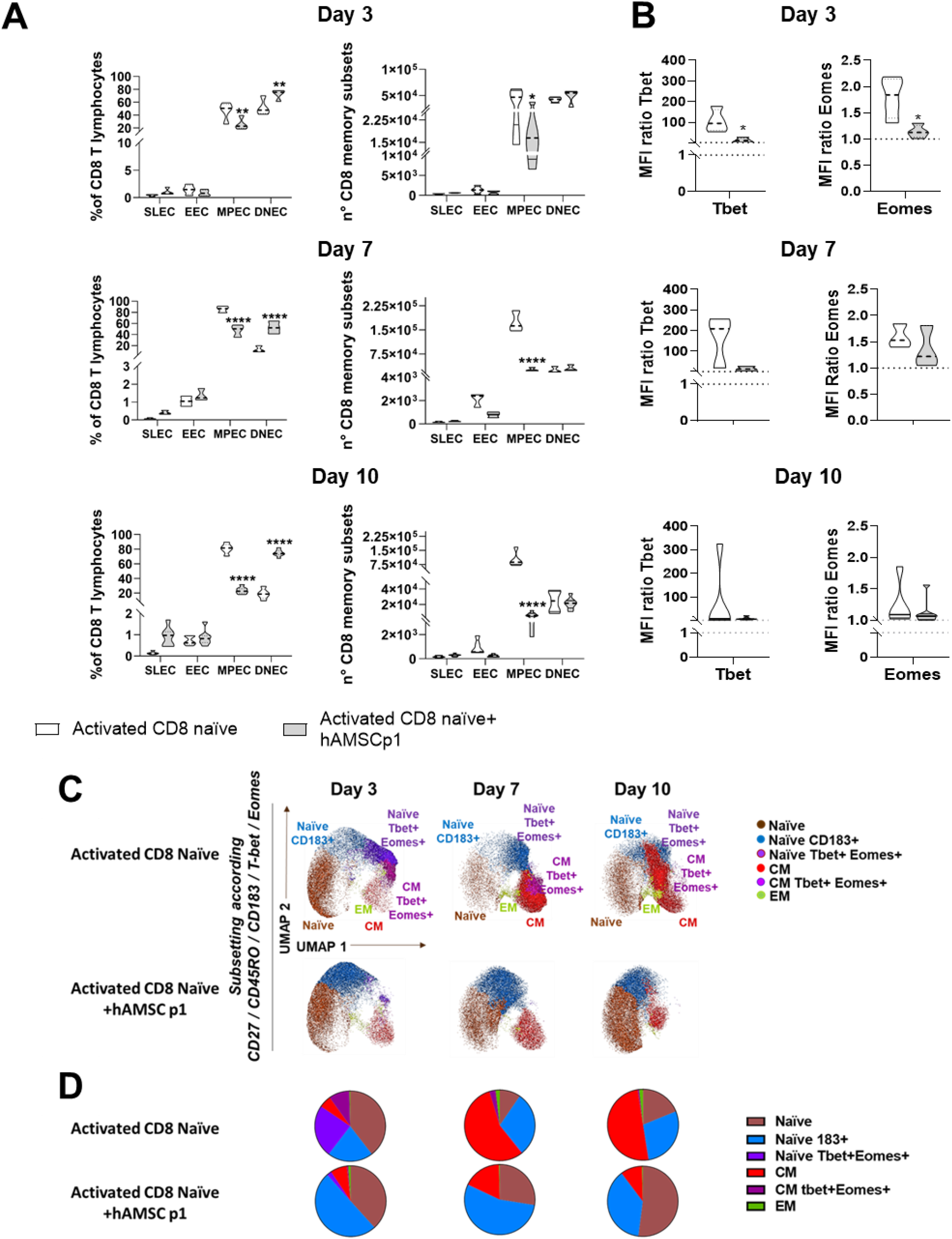
hAMSC influence early phases of T cell commitment. Purified CD8 naïve T lymphocytes were stimulated with antiCD3, antiCD28 and the exogeneous administration of IL12 and IL2, and cultured in the presence of hAMSC (gray). CD8 T cells were allowed to differentiate for 10 days and the different cellular subtypes analyzed at day 3, 7 and 10.(A) CD3+CD8+ T lymphocytes were further analyzed for the expression of CD183 and KLRG1 and classified into four populations: SLECs (CD183-KLRG1+), EECs (CD183+ KLRG1+), MPECs (CD183+ KLRG1-) and DNECs (CD183-KLRG1-). (B) The expression of the transcription factors Tbet and Eomes was evaluated at all timepoints. (C) Uniform Manifold Approximation and Projection (UMAP) representation of the CD8 T cell landscape obtained by Clusterexplorer plugin.(D) Pie charts representing the distribution levels of the 6 identified clusters as a percentage of the total CD8 pool. Results in (A) and (B) are displayed as violin plots showing median (dashed line), 25th and 75th quartiles (*p < 0.05, **p < 0.01, ****p < 0.001); activated CD8 T cells alone represent the control group (white); N ≥ 3 independent experiments.

We next sought to identify the appearance of new CD8 T cell subsets or clusters over time. The unsupervised analysis was performed as aforementioned and shown in Fig. 3C. Co-culture with hAMSC allowed to retain a large amount of naïve T lymphocytes (in brown) and increased the proportion of CD183 naïve T lymphocytes (blue), suggesting that hAMSC do not affect the early stages of activation of naïve CD8 lymphocytes. However, hAMSC strongly reduced the percentage of naïve T lymphocytes positive for the expression of Tbet and Eomes (purple) as well as the commitment towards the CM (red and dark purple) and EM (green) memory subsets (Fig. 3D) hAMSC affect T cell metabolism The PI3K/mTOR pathway is involved in the activation, proliferation, and differentiation of CD8 T lymphocytes (*28*). Because mTOR and its two main complexes, mTORC1 and mTORC2, control the differential expression of Tbet and Eomes (*29, 30*), we next wanted to investigate the effect of hAMSC on these important regulators of cellular metabolism. After three days of co-culture with hAMSC, the total and phosphorylated forms of AKT and mTOR were reduced in activated CD8 T lymphocytes (Fig. 4A). These observations suggest that hAMSC may negatively affect the metabolic switch of naïve CD8 T lymphocytes from fatty acid to glycolytic metabolism, which is required for full maturation of CD8 T lymphocytes (*31*). Representative estimation plots of the MFI ratio between total protein and relative phosphorylated protein show that hAMSCs reduce the phosphorylation of AKT, resulting in a strong reduction in mTOR phosphorylation and activation (Fig. 4B). This suggests a possible mechanism underlying the function of hAMSC to counteract the activation and differentiation of naïve CD8 lymphocytes by inhibiting AKT and mTOR signaling.

**Fig. 4.**
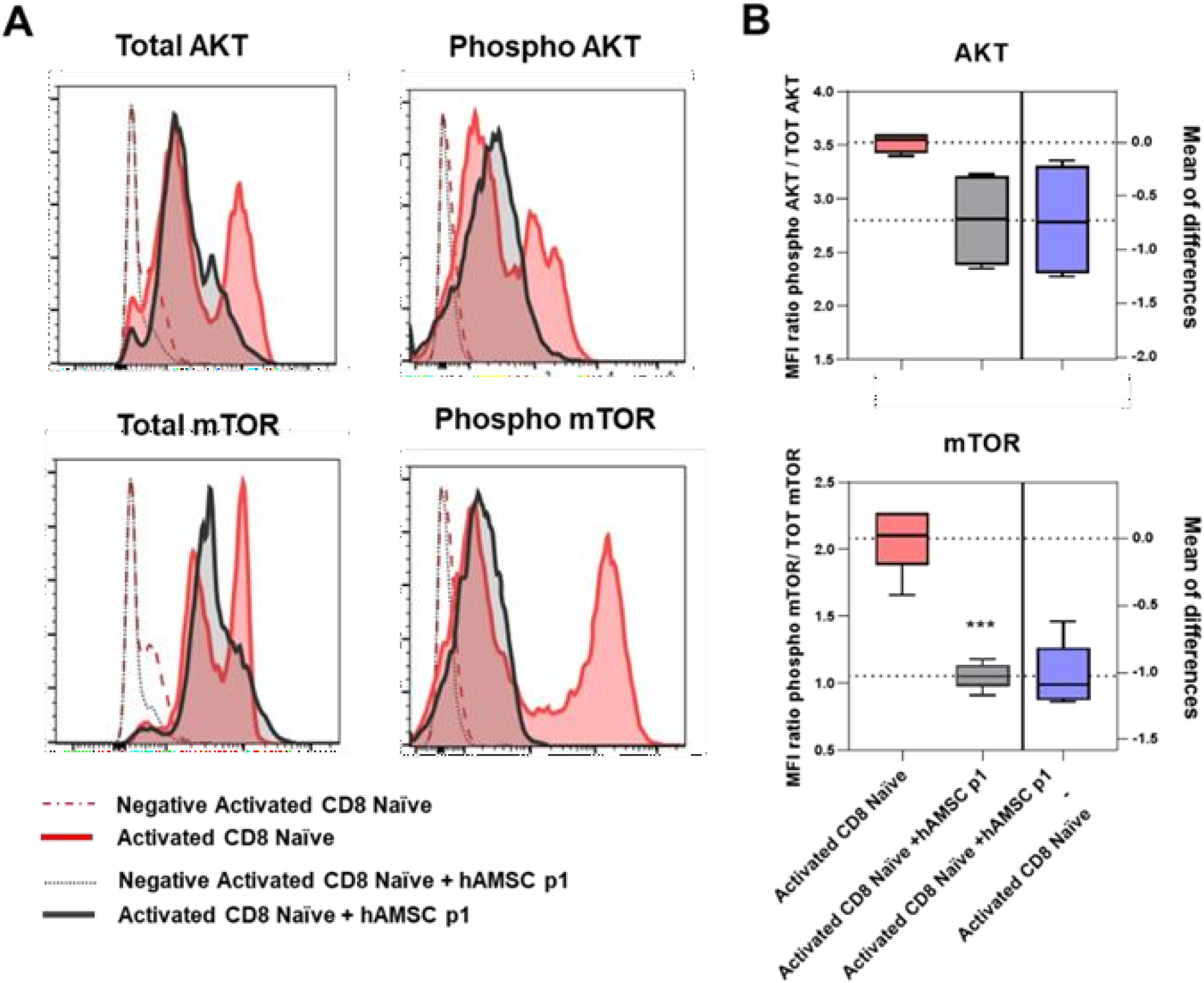
Reduction of AKT and mTOR phosphorylation upon co-culture with hAMSC. Purified CD8 naïve T lymphocytes were stimulated with antiCD3, antiCD28 and the exogeneous administration of IL12 and IL2, and cultured in the presence of hAMSC. Cells were harvested at day 3 and analyzed by flow cytometry. Specifically, CD8 T cells were stained for phospho mTOR, phospho AKT and for their total protein content. (A-B) Representative plots of the MFI ratio between total protein and relative phosphorylated protein. ***p < 0.001; activated CD8 T cells alone represent the control group; N ≥ 3 independent experiments.

### hAMSC modulate the transcriptional landscape of naïve CD8 T lymphocytes

To outline the transcriptional landscape peculiar to the commitment of naïve CD8 T lymphocytes upon hAMSC stimuli, we performed a high throughput gene expression analysis focusing on several genes involved in differentiation and metabolic processes, as well as epigenetic and transcriptional regulation of naïve CD8 lymphocytes. We performed an unsupervised hierarchical clustering analysis and identified genes whose expression is over two-fold higher and statistically significant (p<0.05) in the early (24h) post-activation stages (Supplementary Fig. 1).

Having confirmed the up-regulation of genes such as IL-12 receptor beta 1 (*IL12RB1*) and interleukin-2 receptor alpha (*IL2RA*), involved in the early stages of CD8 T cell activation, we next interrogated on determinants of cell fate. As shown in Fig. 5, the two main inflammatory cytokines, *IFNy* and *TNFα*, typically induced in the early differentiation steps, were up-regulated in activated cells (*23, 32*). At the same time, also *IRF4* and the two master regulators of SLEC and MPEC commitment, *TBX21* (Tbet) and *EOMES* (Eomesodermin), respectively, were up regulated. Conversely, the expression of STAT genes (*STAT3, 4* and *5*), modulators of both activation and commitment of CD8 T lymphocytes, was higher in naïve CD8 T cells compared to activated cells. Interestingly, when naïve CD8 T lymphocytes were activated in the presence of hAMSC, their gene expression profile was comparable to that of unstimulated naïve CD8 T lymphocytes, with the exception of *IL2RA* and *IRF4*, whose increase upon activation was maintained also in the presence of hAMSC.

**Fig. 5.**
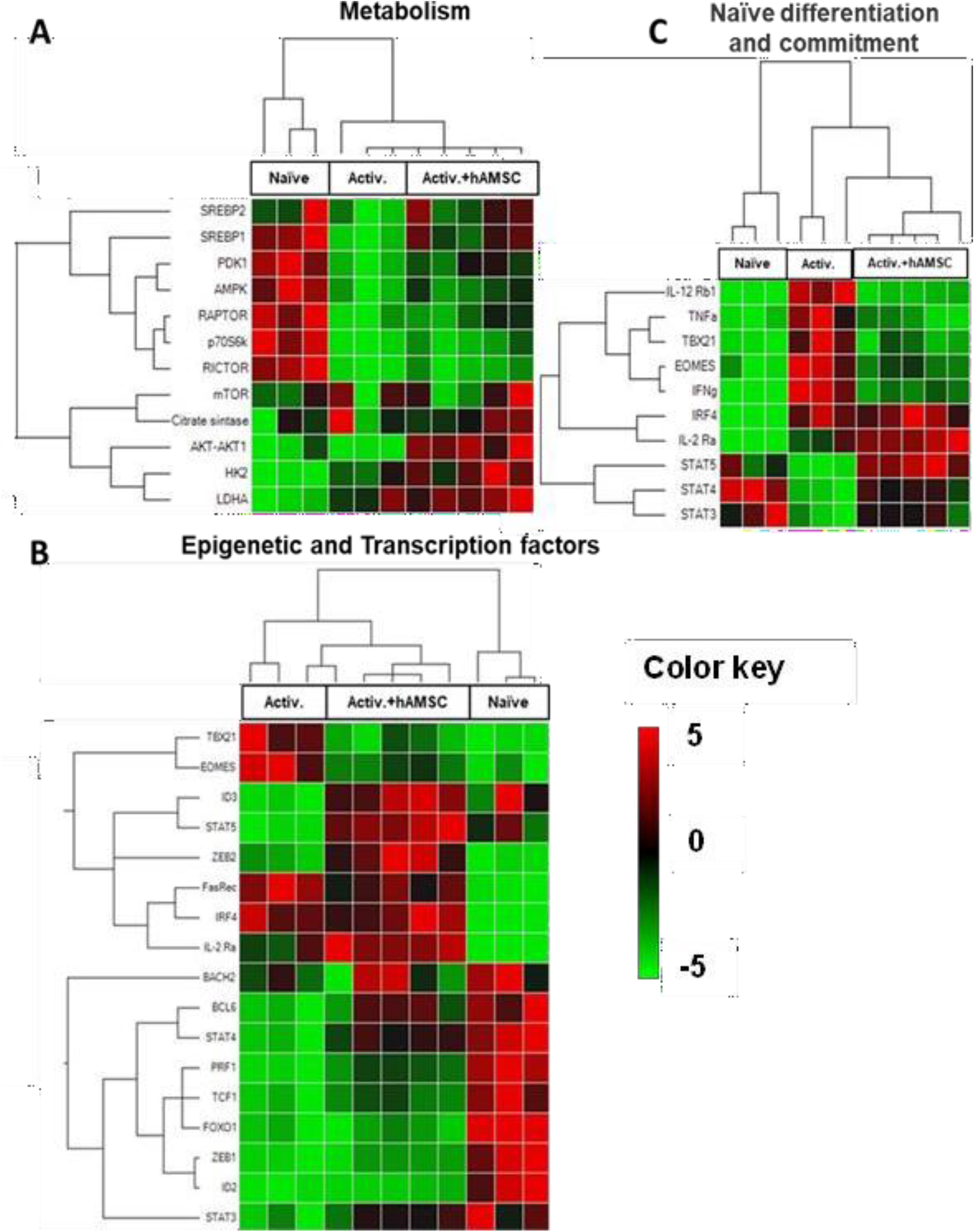
Heatmap chart of gene expression data for cluster analysis. Transcriptomic profile of naïve, activated and activated in the presence of hAMSC CD8 T cells 24 hours upon activation. Panels show metabolic related genes (A), epigenetic and transcriptional factors (B), and genes strictly related to naïve CD8 T cell differentiation (C). The mRNA expression profile is represented as heatmap graph. The genes were clustered according to their expression patterns. N ≥ 3 independent experiments.

Next, we focused our attention on genes involved in metabolic switching. As shown in Fig. 5A, *SREBP1* and *2, AMPK, PDK1, RAPTOR* and *RICTOR* resulted downregulated after stimulation of naïve CD8 T lymphocytes. Again, when naïve CD8 T lymphocytes were activated in the presence of hAMSC, their gene expression profile, and in particular the modulation of *SREBP1* and *2, PDK1*, and *RAPTOR* genes, was comparable to that of unstimulated naïve CD8 T lymphocytes. Similarly, the expression of genes involved in the epigenetic response such as *BACH2* and *ID3* or the transcription factor *BCL6* (*33, 34*) was downregulated after activation and maintained in the presence of hAMSC (Fig. 5B).

Finally, citrate synthase *(CS)*, hexokinase 2 (*HK2*), and *AKT* resulted low expressed in both unstimulated naïve and activated CD8 T lymphocytes, while upregulated upon co-culture with hAMSC.

Overall, our data suggested that hAMSC are not able to completely block CD8 T lymphocytes at the initial naïve stage, but at an intermediate level of differentiation.

### hAMSC regulate the downstream signaling cascade of IL-12 and IL-2 receptor

Given the transcriptional downregulation of IL12Rβ1 on CD8 T cells after hAMSC challenge (Fig. 5C), we speculated on the possible involvement of this receptor in the mechanism of action of hAMSC. To confirm our hypothesis, we examined the expression of the IL-2Rα and that of IL-12Rβ1 at day 3 on both the surface and cytoplasm of activated naïve CD8 T lymphocytes after complete and incomplete stimulation (i.e., in the absence of exogenous IL-12 and IL-2 stimulation). As shown in Fig. 6A, the expression of IL-12Rβ1 on the surface of activated naïve CD8 lymphocytes increased after exogenous stimulation with the two cytokines, but no significant difference in IL-12Rβ1 was observed when CD8 T lymphocytes were cocultured with hAMSC when compared with naïve CD8 T lymphocytes. These results were also confirmed by the mean fluorescence intensity data (Fig. 6B). Instead, a strong decrease in IL-2Rα expression was observed in CD8 lymphocytes cocultured with hAMSC (Fig. 6A-B). Whereas this has been observed previously for IL2-Rα in CD4 (*35*), this result is completely novel for both IL2-Rα and IL-12Rβ1 on CD8 T lymphocytes. Indeed, the expression levels of IL-12Rβ1 in the membrane of naïve CD8 cells cocultured with hAMSC (either incompletely or fully activated) were comparable to those of the control condition (naïve CD8 T lymphocytes activated with the different combination of stimuli) (Figs. 6A-B). This trend was not confirmed at the cytoplasmic level, where co-culture with hAMSC resulted in a decrease in IL-12Rβ1 expression. These results are functionally reflected in a reduction in downstream signaling of the two receptors.

**Fig. 6.**
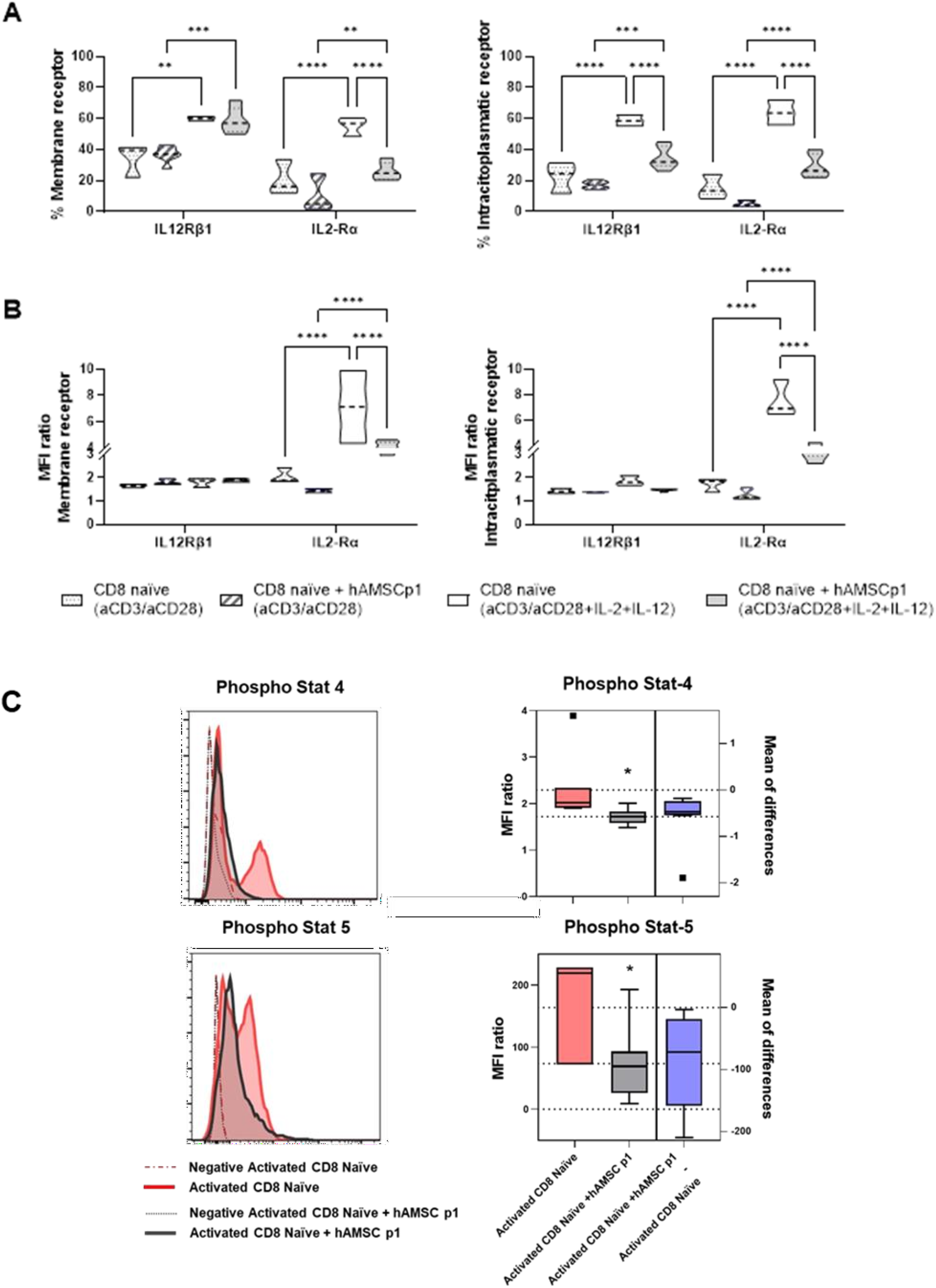
hAMSC regulate the downstream signaling cascade of IL-12 and IL-2 receptor. Purified naïve CD8 T lymphocytes were either stimulated with anti CD3, anti CD28 and by the exogeneous administration of IL12 and IL2 or without the exogenous administration of the two cytokines (not fully activated) in the presence or not of hAMSC.The total amount of IL-12Rβ1 and IL-2Rα expressed on the membrane and in the cytoplasm of CD8 T cells (A) and the level of mean fluorescence intensity (B) were assessed by flow cytometry. The phosphorylation status of the STAT4 and STAT5 was assessed by flow cytometry on fully activated naïve CD8 T lymphocytes co-cultured or not with hAMSC (C). (*p < 0.05, **p < 0.01, ****p < 0.001); N ≥ 3 independent experiments.

Indeed, activation of naïve CD8 T cells in the presence of hAMSC reduced phosphorylation of STAT4 (downstream the IL-12Rβ1 receptor) and STAT5 (downstream the IL2-Rα receptor).

## Discussion

This study sheds new light on the mechanism of action of MSCs isolated from the amniotic membrane (hAMSCs), by explaining for the first time how they affect CD8 T lymphocyte activation and differentiation and their commitment toward memory subsets. Specifically, our study show that hAMSCs block CD8 T cell differentiation and that this is related on the ability of hAMSCs to modulate the expression and downstream signaling of IL-12 and IL-2, receptors which are essential for the full activation and subsequent maturation of naive CD8 lymphocytes (24).

Remarkably, we observed that hAMSCs do not counteract the activation of CD8 T lymphocytes, because after stimulation naïve CD8 T lymphocytes are equally capable of expressing CD183, a marker that is rapidly induced on naïve T lymphocytes after activation (25), but rather decrease commitment to the various memory subsets (CM, EM). This effect has been reported previously for both hAMSCs (11) and bone marrow-derived MSC, although in the latter it was observed only after stimulation with the peptide HY (*36*).

It is well known that cell activation and metabolic activity are closely related. Therefore, we investigated whether hAMSCs are able to modulate the PI3K pathway and consequently the activation of mTOR. The PI3K pathway is involved not only in the activation of T lymphocytes (28), but also in the regulation of metabolism (*37*), and mTOR is considered a master regulator of memory CD8+ T cell differentiation (*38, 39*). We observed that hAMSC are able to decrease the phosphorylation levels of AKT and mTOR, suggesting a possible mechanism to modulate the activation of CD8 T lymphocytes. However, this result is in contradiction with the results of transcriptional analysis, where we observed increased expression of AKT, mTOR, LDHA, and citrate synthase, all genes whose expression is directly involved in the regulation of the various metabolic pathways underlying T-cell activation and early commitment (*40*). These findings suggest that hAMSC do not completely block activation triggered by TCR and costimulatory molecules, but rather interfere with subsequent differentiation processes. By analyzing the expression of the two receptors involved in CD8 memory commitment, we observed that hAMSCs downregulate the expression of IL12RB1 transcript while upregulating the expression of IL2RA.

Analysis of membrane receptors, however, gave us unexpected results. While we expected a decrease in phosphorylation of STAT5 as a consequence of translational blockade of IL2RA, the similarity of the levels of IL-12Rβ1 in naïve CD8 T cells with or without hAMSC conditioning suggests that there is no difference in STAT4 phosphorylation. Interestingly, we observed that STAT4 phosphorylation was reduced when naïve CD8 T lymphocytes were co-cultured with hAMSC.

This finding leads us to hypothesize that hAMSC either induce a phenotype of CD8 T cells that is less responsive to exogenous stimulation or a potential blockade of epigenetic or cross-mediated activation pathways. Another possible explanation is related to the potential alteration in the expression of IL-12Rβ2 whose heterodimerization with IL-12Rβ1 is required for full activation of the receptor and subsequent phosphorylation of STAT4 (*41*). Another possible target of hAMSC activity could be the IL-23 receptor, which also heterodimerizes with IL-12Rβ1, leading to dual phosphorylation of STAT4 and STAT3 downstream of IL-12Rβ1 and IL-23, respectively (*41*). Moreover, hAMSC may result in a non-fully functional IL -12Rβ1 activation, as previously reported (*42*), which can explain the observed reduction in signal transduction consequently impacting on the phosphorylation of STAT4.

Furthermore, the lack of IL-2Rα translation means that ligands, even when administered exogenously, do not trigger the transduction cascade, which in turn could explain the lower activation induced by hAMSC.

The low activation status is also reflected in the lack of IFNγ and TNFα expression. Indeed, the IFNγ promoter is highly methylated in naïve CD8 lymphocytes (*43*). hAMSC likely retain this property in naïve CD8 T lymphocytes, suggesting epigenetic regulation by hAMSC that correlates with their immunomodulatory capacity.

Differentiation of naïve CD8 lymphocytes is regulated by a dynamic and tightly coordinated process involving Tbet and Eomes (*23, 24, 44*) as well as other transcription factors such as IRF4, BCL6, and epigenetic factors such as BACH2, and ID3 (*27, 45, 46*). We observed that hAMSCs modulate this innate cellular commitment and, in particular, affect T lymphocyte activation and differentiation into SLEC and MPEC subsets. TBX21 and EOMES, the two major regulators of SLEC/MPEC fate, are transcriptionally downregulated by hAMSC as in nonactivated naïve CD8 T cells. Consistently, protein expression of Tbet and Eomes was also decreased, which may be related to the observed reduction in STAT4 and STAT5 phosphorylation (*47, 48*).

Furthermore, hAMSC are able to affect the expression of BACH2 by decreasing the availability of the AP -1 complex at transcription sites thus inducing the repression of genes related to terminal differentiation of T lymphocytes (*49*).

In addition, we observed that hAMSCs modulate the expression of ID2 and ID3, two genes that have been reported to be involved in the regulation of SLEC or MPEC commitment (*33*). IL-2, IL-12 and IL-21 have been described to upregulate ID2 and downregulate ID3 expression. We show that hAMSCs impact the expression of ID2 and ID3 by reducing IL-2Rα and IL-12Rβ1 signal transduction, thus explaining the high level of ID3 expression comparable to that of naïve CD8 T lymphocytes, and at the same time the reduction of ID2 expression. Conversely, ZEB2, a target of Tbet normally involved in differentiation processes and induction of effector CD8 T lymphocyte differentiation (*50*), was upregulated by co-culture with hAMSC.

In conclusion, our work unravels a novel immunological mechanism of hAMSC that controls the transcriptional profile of CD8 T lymphocytes between that of naïve and activated cells, thereby influencing the formation of an immunological memory. One of the major implications of our study is that for the first time it focuses on a poorly-studied aspect that is mostly neglected in the field of MSC, namely the effects of MSC on CD8 lymphocytes. Although MSC remain one of the most studied cell therapies for the treatment of immune-related diseases due to their immunoregulatory capabilities, it remains critical to understand what will occur in a patient’s immune system and if the treatment may have side effects. In addition, we report for the first time a mechanism of action for hAMSC to explain the observed effect.

Moreover, it supports the possibility of administering multiple doses of MSC, even from different donors, and highlights their therapeutic potential in graft-versus-host disease (*6*) or autoimmune diseases in which CD8 T lymphocytes play an important role (*51, 52*).

## Materials and methods

### Ethics statements

The collection of human peripheral blood mononuclear cells (PBMC) for research purposes was approved by the Regional Departments of Transfusion Medicine (Rif. 523, July 7, 2016). PBMC were obtained from healthy adult donors after informed consent and provided by Center of Immune Transfusion of Spedali Civili of Brescia, Italy.

Human term placentae were collected from healthy women after vaginal delivery or caesarean section at term, after obtaining informed written consent, according to the guidelines set by the local ethical committee “Comitato Etico Provinciale di Brescia,” Italy (number NP 2243, January 19, 2016).

### Isolation and culture of Human Amniotic Mesenchymal Stromal Cells

Placentas were processed immediately after collection and human amniotic mesenchymal stromal cells (hAMSC) isolated as previously described (*12*) Then the cells were plated and expanded until passage 1 (hAMSC p1) at a density of 1 x104 /cm2 in Chang medium C (Irvine Scientific, Santa Ana, CA, USA) supplemented with 2 mM L-glutamine at 37 °C in the incubator at 5% CO2. Upon reaching confluence, adherent cells were trypsinized and frozen in fetal bovine serum (FBS, Merck, St. Louis, MO, USA) with 10% DMSO (Merck) and stored in liquid nitrogen. hAMSC p1 were phenotypically characterized as previously reported (12). Cells that had > 98% expression of mesenchymal markers CD13 and CD90, < 2% of hematopoietic marker CD45 and < 2% of epithelial marker CD324 were used in this study.

### Isolation of PBMC and naïve CD8 T lymphocytes

PBMC were separated from Buffy Coat through density gradient centrifugation (Histopaque, Sigma-Aldrich,

St. Louis, MO, USA), then frozen in FBS with 10% DMSO (Merck) and stored in liquid nitrogen. Naïve CD8 T lymphocytes were purified from total PBMC by Naïve CD8+ T Cell Isolation Kit and MACS® separation columns (Miltenyi Biotec, Bergisch Gladbach, Germany), following manufacturer’s instructions. Naïve CD8+ T lymphocyte was analyzed with flow cytometry by CD3, CD8, CD197/CCR7, CD45RO expression and the purity resulted >95-98% of the total cells recovered. For the molecular analysis 1 × 106 Naïve CD8 lymphocytes were centrifuged and pellets were stored at -80°C before the RNA extraction.

### Activation of T Cells in PBMC and co-culture with hAMSC

PBMC (1 × 105/well in a 96-well-plate) were seeded in Ultraculture complete medium, composed of Ultraculture medium (Lonza, Basel, Switzerland), supplemented with 2 mM L-glutamine and 1% P/S (both from Sigma-Aldrich) and they were activated with 125 ng/mL (final concentration) anti-CD3 (clone OKT3, BD Biosciences, Franklin Lakes, New Jersey, USA).

hAMSC p1 were plated in RPMI complete medium (composed of RPMI 1640 medium supplemented with 10% heat-inactivated fetal bovine serum (FBS), 1% penicillin and streptomycin, and 1% Lglutamine (all from Euroclone, Pero, MI, Italy), left to adhere overnight (O/N) and γ-irradiated at 30Gy to block their proliferation. Activated PBMC (PBMC + anti-CD3) were cultured in the presence or absence (control condition) of 1 × 105 hAMSC p1. Flow cytometry analysis was performed at day 3 and 7.

### Activation and differentiation of naïve CD8 T lymphocytes and co-culture with hAMSC

Naïve CD8 T lymphocytes (1 × 105/well in a 96-well-plate, and 1 × 106/well in a 24-well-plate) were seeded in Ultraculture complete medium and partially activated with dynabeads human T-activator CD3/CD28 (final dilution 1:1000) (Thermo Fisher Scientific, Waltham, MA USA) or completely activated with dynabeads human T-activator CD3/CD28 and IL2 2.5 U/ml, IL12 10 ng/ml (both from Miltenyi Biotec, Bergisch Gladbach, Germany).

hAMSC p1 were plated in RPMI complete medium, left to adhere O/N and γ-irradiated at 30Gy. Naïve CD8 T lymphocytes (complete or partially activated) were cultured in the presence or absence (control condition) of hAMSC p1 (1 × 105/well in a 96-well-plate, and 1 × 106/well in a 24-well-plate). Flow cytometry analysis was performed at day 3, 7 and 10. For molecular analysis, cells were collected at day 1 and CD8+ T cells were purified from hAMSC by positive selection using CD8 microbeads and MACS® separation columns (Miltenyi) according to the manufacturer’s instructions. After separation, the cells were centrifuged and pellets were stored at -80°C for RNA extraction.

### Flow Cytometry Analysis of CD8 T lymphocyte polarization

Cells were collected at day 3 and 7 (PBMC) and at day 3,7 and 10 (naïve CD8 T lymphocytes), and dead cells were excluded using the eBioscienceTM Fixable Viability Dye eFluorTM 780 (Thermo Fisher Scientific) according to the manufacturer’s instructions. Cells were subsequently stained for 20 min a 4°C with the appropriate combinations of fluorochrome-conjugated anti-human antibodies for the identification of CD8 T cells and sub-populations: CD3 BV480 (clone UCHT1, BD Biosciences), CD8 BV421 (clone RPA-T8, BD Biosciences), CD197/CCR7 Alexa Fluor® 647 (clone 3D12, BD Biosciences), CD45RO PE-CF594 (clone UCHL1, BD Biosciences), CD27 PerCP-Cy5.5 (clone M-T271, BD Biosciences), CD183/CXCR3 PE (clone 1C6/CXCR3, BD Biosciences), KLRG1 PE-Vio770 (clone REA261, Miltenyi), CD212/IL12Rβ1 BV605 (clone 2.4E6, BD Biosciences) and CD25/IL2RA BUV563 (clone 2A3, BD Biosciences).

After membrane staining, intracellular staining with CD212/IL12Rβ1 and CD25/IL2RA was performed after fixation and permeabilization using BD Cytofix/Cytoperm (BD Biosciences).

Proliferation of the T lymphocytes was assessed with Ki67+ Alexa 488 (clone B56, BD Biosciences) and Tbet APC (clone REA102, Miltenyi) and Eomes FITC (clone WD1928, Invitrogen) were assessed by staining with conjugated monoclonal antibody after intranuclear permeabilization used Transcription Factor buffer kit (BD).

Samples were acquired on FACS Symphony A3 BD. Data were analyzed with FlowJo 10.8.

### Determination of phosphorylation status

Three days after co-culture the cells were fixed by adding 1.5% methanol-free formaldehyde (ThermoFisher) for 10 min at room temperature (RT). Then, the cells were collected and stained for 20 min at 4°C with CD8 BV421 (RPA-T8, BD Biosciences). After cell permeabilization in cold 90% methanol for 2-3 days at -80°C, cells were stained in the dark with total AKT and total mTOR (both of Cell signaling Technology, Danvers, Massachusetts, USA) for 30 min at RT, then the cells were washed in stain buffer, consisting of 0.02% sodium azide and 0.1% bovine serum albumin in PBS (Sigma-Aldrich). Cells were then incubated for 30 min at RT with secondary antibody fluorochrome-conjugated anti-Rabbit Dylight 488 (Vector Laboratories, Newark, CA, USA). Finally, cells were stained for 1h at RT with BD PhosflowTM (BD Bioscience) fluorescently conjugated antibodies against: Stat4 (pY693) Alexa fluor® 488 (clone 38/p-Stat4), Stat5 (pY694) PE-Cy7 (clone 47/Stat5(pY694)), Anti-Akt (pS473) PE-CF594 (clone M89-61), Anti-mTOR (pS2448) Alexa fluor® 647 (clone 021-404).

Samples were acquired on FACS Symphony A3 BD. Data were analyzed with FlowJo 10.8.

### Quantitative Real-Time PCR

Total RNA was extracted using EZ1 RNA cell Mini Kit protocol (Qiagen, Frederick, MD, USA), in a BioRobot EZ1 Advanced XL Workstation. The iScript Advanced cDNA Synthesis Kit for RTqPCR (Biorad, Hercules, California, USA) was used for cDNA synthesis, before quantitative real-time PCR cDNA was pre-amplified with SsoAdvanced PreAmp Supermix (Biorad). Real-time PCR was performed using the Biorad instrument CFX96 Quantitative Real-Time PCR. Data were analyzed with Biorad CFX Maestro 2.2 (Biorad).

**TABLE 1.**
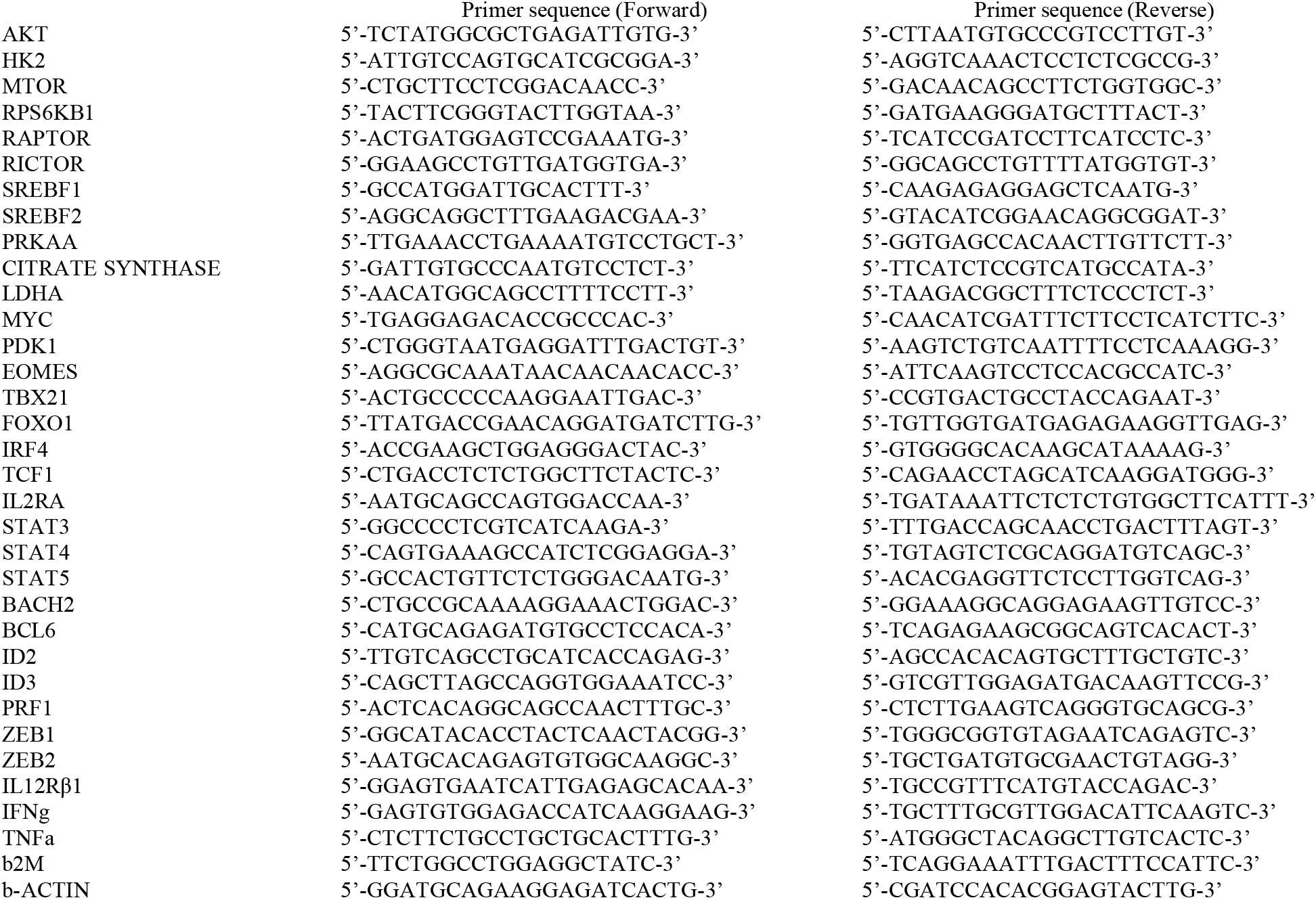
Primer sequences of genes analyzed by quantitative real-time PCR.

## Statistical analysis

Data are shown as violin truncated plots with Tukey variations. The parameters were compared using two-way, one-way analysis of variance (ANOVA) and Student t-test. Data are representative of at least three independent experiments. Statistical analysis was performed using Prism 8 (GraphPad Software, La Jolla, CA, USA). A *p* value lower than 0.05 was considered statistically significant.

## Author contributions

Designing research study: AP, EV and OP. Conducting experiments and acquiring data: EV, PBS, JR. Analyzing data: AP, EV. Result interpretation: AP, EV, PBS, AC, MM. Writing and reviewing the manuscript: AP, EV, AC, MM, FRS, AS, OP. Financial support:AP,OP. Final approval of the manuscript: AP, OP. All authors contributed to the article and approved the submitted version.

## Funding

This work was supported by Ministero della Salute (Ricerca Corrente), Italian Ministry of Research and University (MIUR, 5×1000), PRIN 2017 program of the Italian Ministry of Research and University (MIUR, grant no. 2017RSAFK7), and Contributi per il funzionamento degli Enti privati che svolgono attività di ricerca - C.E.P.R. (2020-2022). Fondi di Ateneo-Linea D1 2019,2020, 2021 (OP), 2022 (AP).

## Acknowledgments

The authors thank the physicians and midwives of the Department of Obstetrics and Gynecology of Fondazione Poliambulanza, Brescia, Italy, and all of the mothers who donated placenta. This work contributes to the COST Action CA17116 International Network for Translating Research on Perinatal Derivatives into Therapeutic Approaches (SPRINT), supported by COST (European Cooperation in Science and Technology).

## Conflict of interest

The authors declare that the research was conducted in the absence of any commercial or financial relationships that could be construed as a potential conflict of interest.

## Caption

**Graphical abstract.**
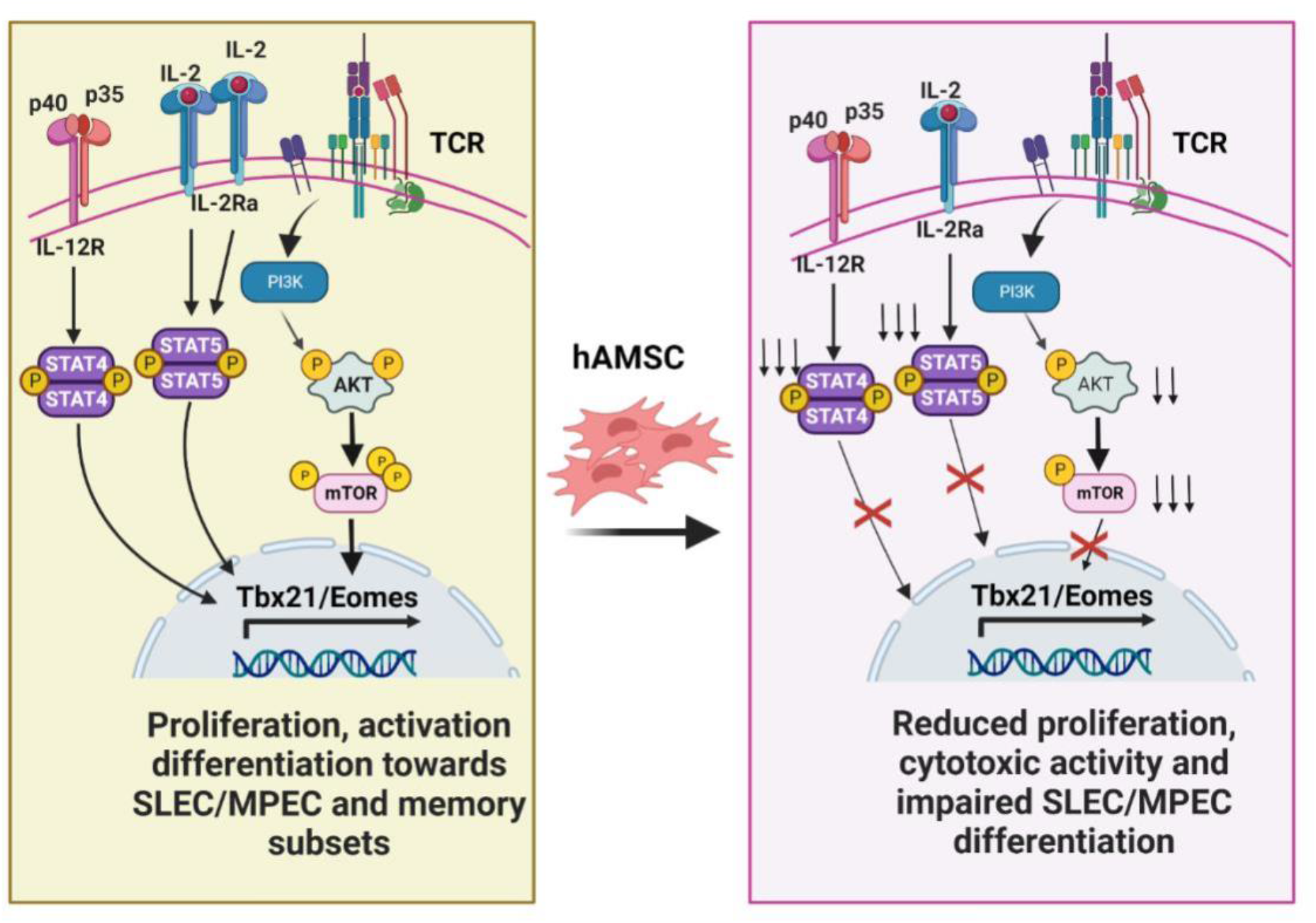
A novel immunological mechanism of hAMSC. hAMSC impair the generation of an immunological memory repertoire by altering the commitment of CD8 T cells towards SLEC and MPEC subsets via Tbet and Eomes. hAMSC reduce the expression of IL-12Rβ1 and IL2Rα receptors, block the phosphorylation of STAT4 and STAT5 downstream and that of AKT and mTOR.

**Supplementary Fig. 1.**
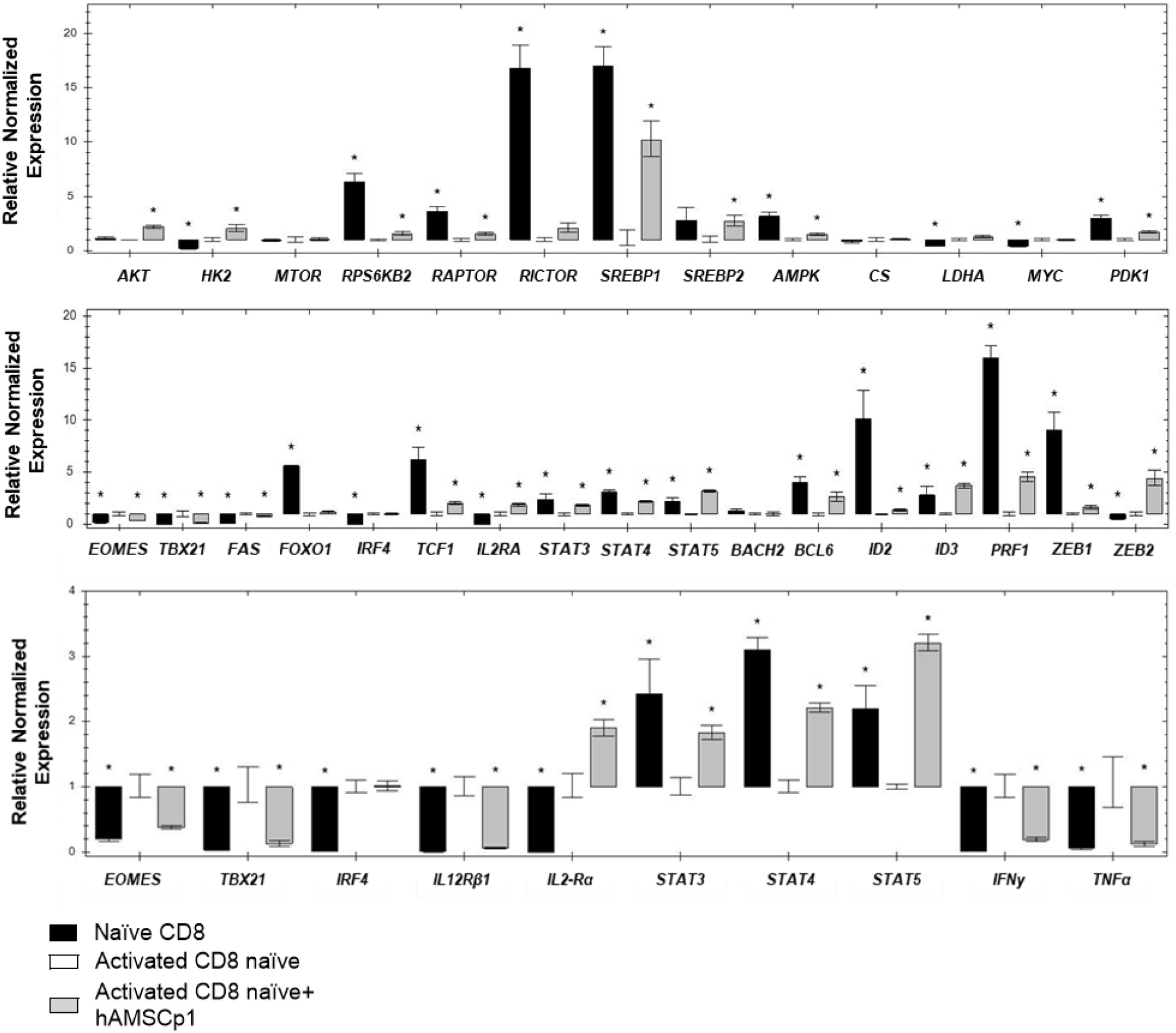
Gene expression analysis. Values shown were normalized on the control condition represented by activated naïve CD8 T lymphocytes. In black the expression value of freshly isolated naïve CD8 T lymphocytes is shown, in gray the gene expression after co-culture with hAMSC. N ≥ 3 independent experiments; (*p < 0.05).

## Notes

### Competing Interest Statement

The authors have declared no competing interest.

### Summary of Updates

Modify affiliation for one of the author

